# Regulatory logic and transposable element dynamics in nematode worm genomes

**DOI:** 10.1101/2024.09.15.613132

**Authors:** Janna L. Fierst, Victoria K. Eggers

**Affiliations:** Biomolecular Sciences Institute and Department of Biological Sciences, Florida International University, 11200 8th Street, 33199, Miami, FL, USA

## Abstract

Genome sequencing has revealed a tremendous diversity of transposable elements (TEs) in eukaryotes but there is little understanding of the evolutionary processes responsible for TE diversity. Non-autonomous TEs have lost the machinery necessary for transposition and rely on closely related autonomous TEs for critical proteins. We studied two mathematical models of TE regulation, one assuming that both autonomous tranposons and their non-autonomous relatives operate under the same regulatory logic, competing for transposition resources, and one assuming that autonomous TEs self-attenuate transposition while non-autonomous transposons continually increase, parasitizing their autonomous relatives. We implemented these models in stochastic simulations and studied how TE regulatory relationships influence transposons and populations. We found that only outcrossing populations evolving with Parasitic TE regulation resulted in stable maintenance of TEs. We tested our model predictions in *Caenorhabditis* genomes by annotating TEs in two focal families, autonomous LINEs and their non-autonomous SINE relatives and the DNA transposon *Mutator*. We found broad variation in autonomous - non-autonomous relationships and rapid mutational decay in the sequences that allow non-autonomous TEs to transpose. Together, our results suggest that individual TE families evolve according to disparate regulatory rules that are relevant in the early, acute stages of TE invasion.

## Introduction

Transposable elements (TEs) have been described as selfish genomic parasites which are dependent on the eukaryotic cell for maintenance and replication (Doolittle and Sapienza, 1980; Orgel and Crick, 1980; Kidwell and Lisch, 2001), but that view has evolved with evidence that TEs can play critical roles in organismal development (Wang et al., 2022; Chang et al., 2024) and gene regulation (Chuong et al., 2016). The emerging picture is that the evolution of TEs can have complex detrimental and beneficial effects for an organism. TEs can rapidly replicate, mutate and insert throughout the genome (Craig et al., 2002), potentially increasing genome size (Pritham, 2009; Tenaillon et al., 2011; Lehmann et al., 2021). Replication can harm genes and essential functions through disruptive insertions (Finnegan, 1992) but also create pools of novel genetic variation and polymorphism (Fedoroff, 2012). TEs can influence selection and molecular evolution (Castillo et al., 2011) and modulate gene expression levels (Hatanaka et al., 2024). The evolutionary processes generating patterns of TE diversity are not well understood and discovering the factors driving TE dynamics is a central question in biology.

Some TEs are clearly molecular parasites. Autonomous transposons produce the machinery necessary for transposition, including intact recognition sequences, competent proteins and critical enzymes such as transposase, integrase and transcriptase (Makalowski et al., 2019; Fig. 1). In contrast, non-autonomous transposons have lost their transposition machinery through mutation (Wicker et al., 2007). They rely on closely related autonomous copies for transposition (McClintock, 1950) potentially parasitizing both the genome itself and their transposon relatives (Hartl et al., 1992).

**Fig. 1.**
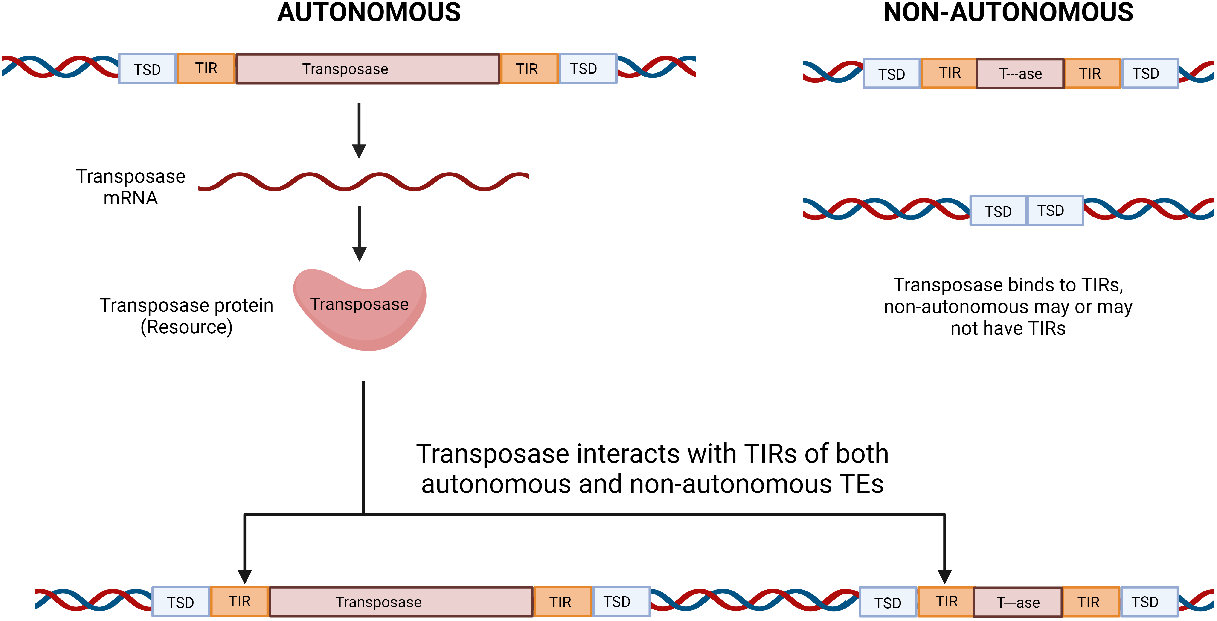
Autonomous TEs are characterized by competent recognition sequences including target site duplications (TSDs) and terminal inverted repeats (TIRs). They also encode full-length enzymes, here transposase. The transposase protein must be shared between autonomous and non-autonomous relatives and becomes a limiting resource in transposition. Non-autonomous TEs have independent and combined deletions of the sequences necessary to produce competent enzymes and TSD and TIR recognition sequences.

Autonomous and non-autonomous transposons do not share the same interests and likely do not obey the same regulatory logic (Hartl et al., 1992). Empirical evidence suggests the two types of TEs transpose and proliferate differently (Hartl et al., 1992; Robillard et al., 2016). TE insertions can damage critical genes and disrupt genome integrity to the point of harming or killing the host (Finnegan, 1992; Charlesworth et al., 1994). The replication of autonomous transposons is limited by their ability to produce proteins and enzymes such as the transposase necessary for successful insertion. For these reasons, autonomous transposons may have a strong interest in attenuating replication and mutational insertions (Doolittle et al., 1984).

Non-autonomous transposons compete with their autonomous relatives for critical enzymes. Capturing transposition machinery for *trans* regulation (McClintock, 1950) puts them at a substantial disadvantage for the higher affinity resources produced in *cis* by autonomous transposons (Hartl et al., 1992; Fig. 1). For non-autonomous transposons, attenuating activity would potentially result in losing the ability to transpose all together.

Here, we ask how transposon regulatory relationships influence evolutionary trajectories by studying two deterministic mathematical models, implementing our models in stochastic simulations, and testing our predictions in nematode genome sequences. Previous models of TE evolution have assumed that autonomous and non-autonomous transposons follow the same regulatory rules (Brookfield, 1996; Le Rouzic and Capy, 2006). In this study we developed two models of TE regulatory logic, one where autonomous elements self-attenuate while non-autonomous elements increase transposition, parasitizing their autonomous relatives (Fig. 2A) and one where both autonomous and non-autonomous elements continually increase transposition rates and compete (Fig. 2B).

**Fig. 2.**
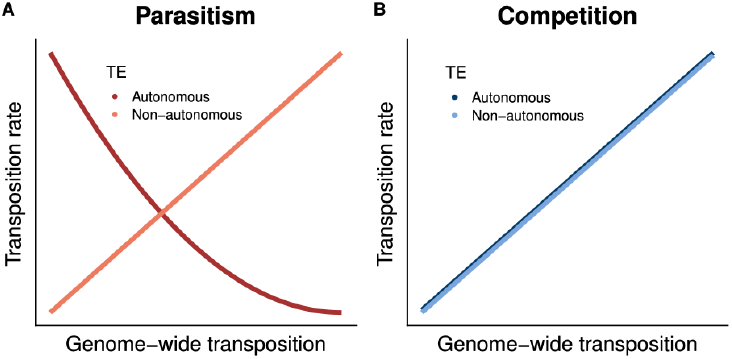
Under A) a parasitic model of regulatory logic autonomous TEs self-attenuate transposition while their non-autonomous relatives continue to increase transposition rates as genome-wide transposition increases. Under B) a competitive model of regulatory logic both autonomous and non-autonomous transposons increase transposition rates as genome-wide transposition increases. Both autonomous and non-autonomous TEs share the same function but are plotted here with a small displacement for visualization.

TEs are subject to stochastic factors in real populations, some resulting in loss due to drift. To study these processes we implemented our deterministic equations in stochastic, individual-based population genetic simulations and studied the evolution of populations with differing modes of reproduction. Self-fertility and outcrossing influence population genetic factors including the effective population size, recombination rate, genetic variation and ultimately the probability a TE is lost or establishes in the population. Previous models have predicted extreme effects of sexual and asexual reproduction ranging from extinction of the TE in the genome in asexual populations (Hickey, 1982) to population extinction in both sexual (Charlesworth and Langley, 1986) and asexual species (Arkhipova and Meselson, 2005). Critically, none of these early theoretical treatments explain the patterns of TE proliferation and diversity we see in genomes today.

Our deterministic models predicted stable coexistence of both TE types but under population genetic constraints and stochastic conditions only large sexually outcrossing populations evolving with Parasitic regulatory logic maintained stable numbers of autonomous and non-autonomous TEs. The stochastic model of Competitive TE regulatory logic predicted autonomous TEs outnumber their non-autonomous counterparts while the stochastic model of Parasitic TE regulatory logic predicted the opposite pattern with non-autonomous TEs outnumbering autonomous. We tested these predictions by examining two TE superfamilies containing both autonomous -non-autonomous members, the retrotransposons LINEs/SINEs and the DNA transposon*Mutator*, annotated in assembled *Caenorhabditis* genome sequences. We found that autonomous LINEs far outnumber non-autonomous SINEs, supporting a model of Competitive regulation. In contrast, non-autonomous *Mutator* TEs far outnumbered autonomous elements, supporting a model of Parasitic regulation.

Non-autonomous *Mutator* TEs require competent terminal inverted repeat (TIR) sequences for transposase binding. We found that in *Caenorhabditis* these sequences rapidly mutate beyond recognition, suggesting non-autonomous TEs are quickly rendered transposition-incompetent. Our theoretical results demonstrate that regulatory relationships can create intrinsically stable or ephemeral systems of TEs, and our comparative genomic results suggest broad variation in the regulatory logic underlying the evolution of individual TE families. Rapid mutational decay limits the influence of regulatory logic to early stages of invasion, with inactivated sequences lingering in the genome as relics of past TE dynamics.

## Methods

### The deterministic model

We developed two deterministic models of transposon dynamics, building on previous analytical work (Charlesworth and Charlesworth, 1983; Dolgin and Charlesworth, 2006, 2008; Le Rouzic and Capy, 2006). Both models envision two types of TEs, one fully autonomous and capable of producing all machinery necessary for transposition and one non-autonomous, dependent on its autonomous relative for the sequences and enzymes necessary for transposition. Autonomous element *i* and non-autonomous element *j*, respectively, have transposition activities *a*_*i*_ and *a*_*j*_ , roughly the probability of transposition per element. If *n*_*i*_ is the number of autonomous elements and *n*_*j*_ is the number of non-autonomous elements, then *n* is the total of all TEs in an individual. Each TE carries a small additive fitness cost *s* and the mean fitness *w*_*n*_ of an individual with *n* TEs is a linear function, *w*_*n*_ = 1 + *s ∗ n*.

TEs insert into the genome according to *u*_*i*_, the transposition ability of TE *i* and excise out of the genome according to *ν*_*i*_. Here, autonomous and non-autonomous tranposition abilities are equal and *u*_*i*_ = *u*_*j*_ while *ν*_*i*_ = *ν*_*j*_ . The mean transposition activity for the genome, 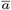, encompasses the transposition of all autonomous and non-autonomous TEs. Under a parasitic model of TE regulation, the autonomous TE *i* attenuates transposition with increasing genome-wide tranposition activity and its transposition rate is given by 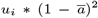 (Fig. 2B). In contrast, the non-autonomous TE *j* does not attenuate as transposition activity increases, transposing instead according to 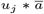. Under a competitive model of TE regulation both TEs transpose according to 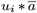, continually amplifying competition for physical transposition resources (Fig. 2A).

Mean transposition is calculated as

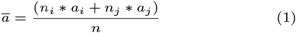

and the rate of change in the population abundance of transposons is determined by

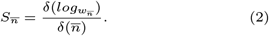

At low copy number TE frequencies can be approximated by a Poisson distribution (Le Rouzic and Capy, 2006) where 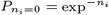 and 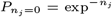. The change in TE copy number between generations in the parasitic model of regulation is then calculated according to

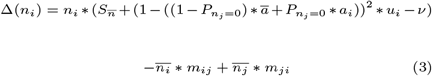

for autonomous TEs and

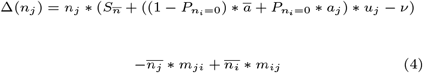

for non-autonomous TEs. The terms *m*_*ij*_ and *m*_*ji*_ represent the potential for elements to mutate from autonomous to non-autonomous and vice versa.

The competitive model of TE regulation evolves according to:

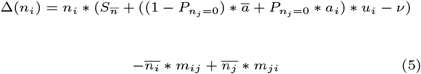

and

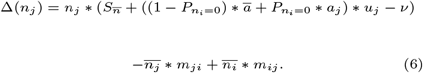

### The stochastic model

TEs are subject to multiple interacting stochastic factors that are difficult to model with deterministic approaches. We implemented our deterministic mathematical models of parasitic and competitive regulation in forward time simulations using SLiM 4.0.1 (Haller and Messer, 2019, 2023) to study the influence of population genetic factors on TE evolution.

*Caenorhabditis* nematodes have two primary mating systems. The ancestral reproductive mode is dioecy with males and females. Three species (*C. elegans, C. briggsae and C. tropicalis*) independently evolved an androdioecious reproductive mode where hermaphrodites produce young by either self-fertilizing or outcrossing with males (Kiontke and Fitch, 2005; Kiontke et al., 2004). Male frequency and outcrossing vary quantitatively from approximately 1 in every 1000 individuals as male in laboratory populations of the *C. elegans* strain N2 to 1 in every 3 individuals as male in populations of the related *C. elegans* strain AB1 (Anderson et al., 2010). Following these features of reproductive mode, our simulated dioecious populations contained males and females. Simulated self-fertile populations had a proportion of individuals reproducing through selfing and a proportion outcrossing. In order to simulate androdioecous populations we constructed ‘non-Wright Fisher’ populations in SLiM 4.0.1 (Haller and Messer, 2019, 2023) with overlapping generations and variable sizes.

We evolved populations under a number of different evolutionary scenarios including small and large population size (ranging between *N* = 100 and *N* = 1000, respectively), high and low recombination rates (ranging between 1*x*10^*−*2^ and 1*x*10^*−*5^) and different levels of self-fertility ranging from full dioecy with 0% of the population reproducing through self-fertility to 100% of the population reproducing through self-fertility. Each individual had a single diploid chromosome 1*x*10^5^. Each population was initialized with 4 TEs (2 on each chromosome) with selective effect *s* = 0.001. TEs inserted at a rate *µ* = 0.01 and excised at *ν* = 0.001. The activity *a*_*i*_ of the autonomous TE was 1 and the activity *a*_*j*_ of the non-autonomous TE was 0. Simulated populations were evolved for 500,000 generations.

### Annotating TEs in genome sequences

We focused on two superfamilies of TEs, one the autonomous retrotransposon Long Interspersed Nuclear Elements (LINEs) and their non-autonomous Short Interspersed Nuclear Elements (SINEs) relatives (Weiner, 2002) and the other autonomous and non-autonomous members of the DNA transposon *Mutator* (Dupeyron et al., 2019; Lisch, 2015; Robertson, 1978, 1986). Although some LINEs have been rendered inactive by mutations (Grimaldi et al., 1984; Ostertag and H H Kazazian, 2001; Beck et al., 2011), members of the family are classified as autonomous and many retain the ability to transpose autonomously. In contrast, SINEs are severely mutated and unable to produce their own transposition machinery. LINEs and SINEs are Class I TEs and require an RNA intermediate to transpose.

In contrast, members of the DNA transposon *Mutator* family are Class II TEs and able to transpose without a molecular intermediate. *Mutator* is a broadly dispersed TE named for its mutagenic potential (Robertson, 1978, 1986). Previous investigations into *Caenorhabditis* genomes revealed large numbers of *Mutator* TEs that are approximately twice as abundant in the genomes of outcrossing species (Adams et al., 2023).

Existing genome assemblies and EDTA annotations were obtained from Adams et al. (2023). Briefly, Extensive *de-novo* TE Annotator (EDTA) v2.0.0 (Ou et al., 2019) annotates TEs by employing several annotation softwares and filtering steps. Long terminal retrotransposons were found by EDTA with LTRharvest (Ellinghaus et al., 2008) and LTR Finder (Ou and Jiang, 2019) before filtering with LTR retriever (Ou and Jiang, 2018). Generic Repeat Finder (Shi and Liang, 2019) identified terminal direct repeats (TDRs), miniature inverted transposable elements (MITEs), and terminal inverted repeats (TIRs). TIRs were also found with TIR-Learner (Su et al., 2019). *Helitrons* were found with HelitronScanner (Xiong et al., 2014). Both TIRs and *Helitrons* were investigated for simple sequence repeats using *MUSTv2* (Ge et al., 2017). Identified TEs were used to mask the genome, and RepeatModeler (Smit et al., 2013-2015) then scanned the unmasked portion to find any remaining TEs. False discovery rate was reduced by supplying EDTA with curated Rhabditida repeat libraries and coding sequences (CDS) to filter the TE library. Curated repeats were obtained from RepeatMasker (Smit et al., 2013-2015). CDS files were generated with BRAKER2 (Bruna et al., 2021) or downloaded from either WormBase Parasite (Howe et al., 2017) or NCBI (National Center for Biotechnology Information, 2002). Finally, EDTA classifies TEs with TEsorter (Zhang et al., 2019).

We assigned TEs as autonomous and non-autonomous through further analysis of EDTA output files (Ou et al., 2019). For each *Caenorhabditis* genome, EDTA files ending in *mod.EDTA.TEanno.gff3 were filtered for a specific TE classification, such as *Mutator*. The first, fourth and fifth columns were cut to make a bed file. We used bedtools getfasta (Quinlan and Hall, 2010) to obtain the TE sequences for each of these specific classification files. Potential genes within individual TE sequences were then identified by searching for open reading frames (ORFs) with a minimum size of 300 base pairs using the function getorf from EMBOSS v.6.6.0 (Rice et al., 2000). These ORFs were supplied to pfam scan.pl v.1.6 to scan PFAM database v.36.0 for functional annotation (Punta et al., 2012). A web scraping python script was created to search for the word “Transposase” on the webpage descriptions of each unique pfamID found in the TE sequences. For LINEs and SINEs “Transposase” was changed to “Integrase” or “Endonuclease,” for example. TE sequences of appropriate length (*>*500 nucleotides for *Mutator* and *>*2000 nucleotides for LINEs) with pfamID descriptions containing these critical proteins were labeled as autonomous while others were labeled as non-autonomous.

For *Mutator*, the DNA binding site for transposase is likely the Terminal Inverted Repeat (TIR) sequences. Thus, parasitism or competition would require non-autonomous elements with TIRs similar enough to be recognized by transposase proteins of autonomous TEs. This was inferred through sequence similarity of TIRs between autonomous and non-autonomous TEs. TIRs were extracted from the last column of EDTA files ending in *mod.EDTA.intact.gff3. Levenshtein distance algorithm was used to compute sequence similarity between TIRs and those with a similarity of at least 80% were classified as active non-autonomous TEs.

## Results

### The deterministic model

When autonomous and non-autonomous TEs have identical rates of transposition *u*_*i*_ and excision *ν*_*i*_, the deterministic model with competitive regulation predicts both elements will achieve the same stable copy number in the populations within 1500 generations (Fig. 3). With parasitic regulatory dynamics the two elements achieve stable copy number in the population within 3500 generations but non-autonomous TEs are more abundant than their autonomous relatives.

**Fig. 3.**
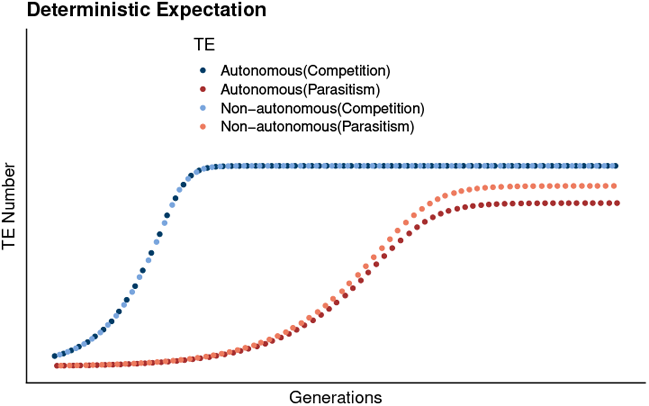
When autonomous and non-autonomous transposons operate according to (A) the same competitive regulatory logic the deterministic model predicts a small number of transposons will increase until they stabilize in the population. The two TEs have identical numerical trajectories and are plotted on top of one another. (B) When the non-autonomous transposon parasitizes their autonomous relative, the transposons stabilize in the population but it takes more than twice as many generations. At a stable equilibrium the non-autonomous transposon is more abundant than the autonomous transposon. Here, *u*_*i*_ = 0.01, *ν*_*i*_ = 0.001 and the selective effect of each TE is *s* = 0.001.

### The stochastic model

We found that the simulated populations followed 4 discrete trajectories according to the population genetic factors they evolved under (Fig. 4). We divided these trajectories into 4 categories to qualitatively describe the impact of population size, recombination rate, reproductive mode and regulatory logic on TE evolution and maintenance. Populations that followed an ‘Early TE Loss’ trajectory would rapidly lose both autonomous and non-autonomous TEs, typically in fewer than 20,000 simulated generations. In other populations both autonomous and non-autonomous TEs were maintained for 20,000-450,000 generations but eventually excised entirely from the population. We labeled populations following this trajectory ‘Eventual TE Loss.’

**Fig. 4.**
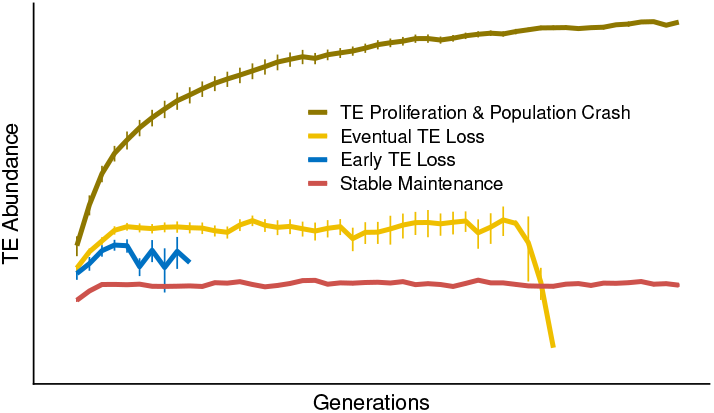
Across the range of population genetic parameters the simulated populations evolved according to 4 trajectories: 1) Early TE Loss; 2) Eventual TE Loss; 3) TE proliferation, fitness loss and eventual population extinction; or 4) Stable Maintenance

Some populations had TEs that continually proliferated and increased in abundance. These populations lost fitness to the point of eventual population extinction. Importantly, some of the simulated populations had unchecked proliferation of both autonomous and non-autonomous TEs while in other populations the non-autonomous TE was lost early on and the autonomous TE subsequently proliferated to the point of population extinction. We labeled both of these scenarios ‘TE Proliferation & Population Crash.’ Finally, some populations maintained both autonomous and non-autonomous TEs over the entire course of the simulated population evolution and we labeled this trajectory ‘Stable Maintenance.’

We found that three population genetic parameters (population size, reproductive mode and regulatory logic) determined the trajectory of TEs in the simulated populations (Table 1). Small populations frequently had trajectories of Early TE Loss across both reproductive modes and models of TE regulation (Supp. Fig. 1). Those populations that maintained TEs followed a trajectory of TE Proliferation & Population Crash with TE insertion increasing to the point at which fitness loss was severe and population-wide. Under a Competitive model of TE regulation some simulated populations decoupled the non-autonomous and autonomous TEs, losing the non-autonomous entirely but maintaining the autonomous TE (Suppl. Fig. 1,2). These populations eventually followed the same TE Proliferation & Population Crash trajectory.

**Table 1.** TE trajectories are determined by population size, reproductive mode and TE regulation. The trajectories are: 1.Early TE Loss 2.Eventual TE Loss 3.TE Proliferation & Population Crash 4.Stable Maintenance

|  |  | Competition |  | Parasitism |  |
| --- | --- | --- | --- | --- | --- |
|  |  | Selfing | Outcrossing | Selfing | Outcrossing |
| Population | Small | 1, 3 | 1, 3 | 1, 3 | 1, 3 |
|  | Large | 1, 2, 3 | 1, 2 | 1, 3 | 4 |

TEs were retained for long periods in some large dioecious populations evolving with Competitive TE regulation (Fig. 5A, B) but gradually lost across the population replicates, following a trajectory of Eventual TE Loss (Fig. 4). TEs were stably maintained in large dioecious populations evolving with Parasitic TE regulation (Fig. 5A, B). Importantly, this was the only set of simulated population parameters that resulted in a trajectory of Stable Maintenance (Table 1; Fig. 3).

**Fig. 5.**
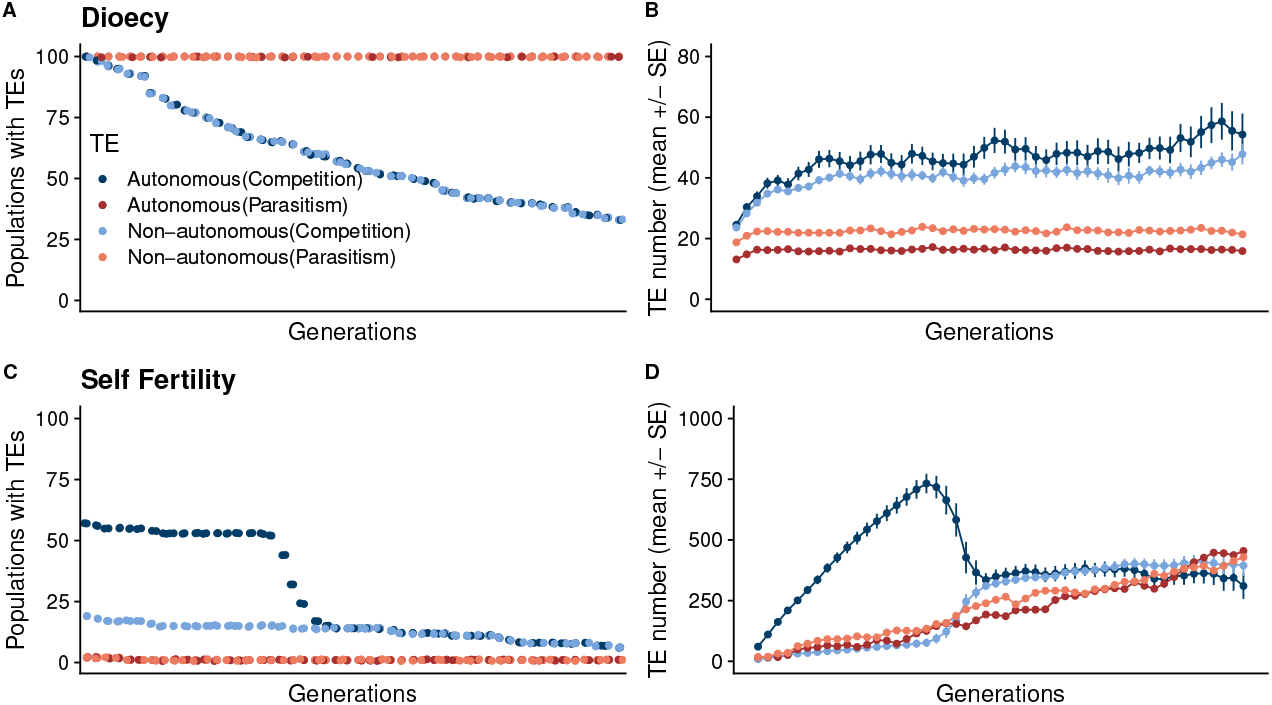
When the population is large (here, N=1000 individuals) and dioecious, populations evolving with (A-B) competitive TE regulation gradually lose TEs. In contrast, populations evolving with (C-D) parasitic TE regulation stably maintain a low number of TEs.

Large self-fertile populations evolving with either Competitive or Parasitic TE regulation followed trajectories of Early TE Loss or TE Proliferation % Population Crash (Table 1; Fig. 5,6). Some large self-fertile populations evolving with Competitive TE regulation decoupled the two TE types, excising the non-autonomous TE entirely and maintaining the autonomous TE, but eventually these populations followed the TE Proliferation % Population Crash trajectory (Fig. 6A-D). At low levels of outcrossing, for example 5% of the population outcrossing and 95% selfing, populations evolving under Competitive or Parasitic TE regulation followed the trajectories of fully self-fertile populations (Fig. 6). At 50% outcrossing populations evolving with either Competitive or Parasitic TE regulation followed the trajectories of fully outcrossing populations (Suppl. Fig. 3).

**Fig. 6.**
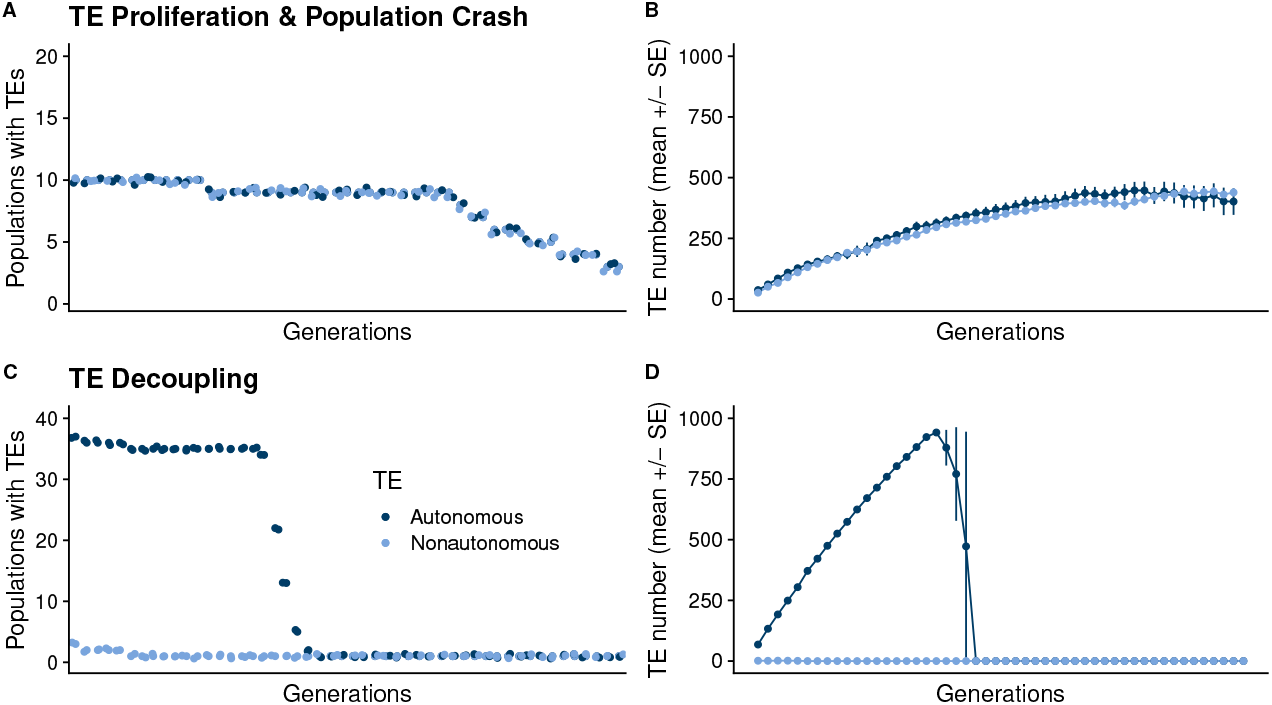
Large self-fertile populations evolving with competitive TE regulation evolved according to different trajectories. (A-B) Ten of the 100 populations retained both TEs for a period of time, eventually losing both autonomous and non-autonomous TEs. (C-D) Thirty-eight of the populations retained their autonomous TE but lost their non-autonomous TE early in the evolution of the simulated populations. Those populations experienced TE proliferation, fitness loss and eventual population crashes.

We quantitatively varied population size and found that population of 500 individuals and smaller followed trajectories similar to population sizes of 100 individuals (Suppl. Fig. 1,4). Population sizes of 750 individuals and larger followed trajectories similar to population sizes of 1000 individuals (Suppl. Fig. 5). Importantly, populations evolving under Competitive TE regulation did not achieve the Stable Maintenance trajectory at large population sizes of 1500 individuals. Similarly, we found that varying recombination rate altered the speed with which the simulated populations achieved different trajectories but did not qualitatively change the trajectories determined by population size, reproductive mode and TE regulation (Suppl. Fig. 2). In population genetic models of TEs the rate of recombination heavily influences linkage disequilibrium among suites of TEs (Roze, 2023) depending on the shape of the fitness function (Charlesworth, 1990).

### Testing predictions from the model

Importantly, the deterministic and stochastic models predict that autonomous and non-autonomous TEs should have differing dynamics in empirical populations. Populations evolving with Competitive TE regulation should have higher abundances of autonomous TEs while populations evolving with Parasitic TE regulation should have higher abundances of non-autonomous TEs. To test these predictions and evaluate the regulatory logic operating in eukaryotic genomes we studied annotated autonomous and non-autonomous TEs in assembled *Caenorhabditis* genome sequences.

We found that LINE elements were more abundant than SINE elements across both androdioecious and dioecious *Caenorhabditis* species (Fig. 7). The number of annotated LINE elements ranged from 149-2258 in dioecious species and 599-1379 in androdioecious species. In comparison, SINE elements ranged from 0-31 in dioecious species and 1-487 in androdioecious species.

**Fig. 7.**
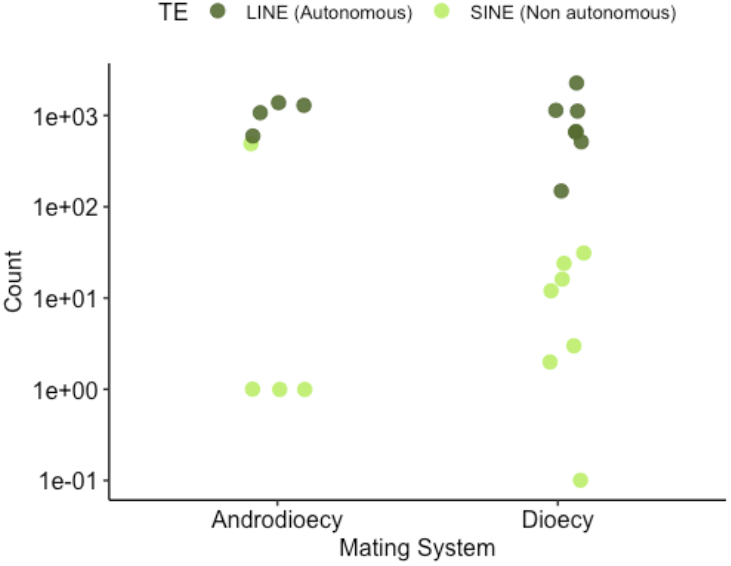
Autonomous LINE elements are more abundant than mutated non-autonomous SINE elements in both androdioecious and dioecious *Caenorhabditis* genomes.

*Mutator* elements ranged from 8,796 to 36,880 in *Caenorhabditis* (Table 2). The number of non-autonomous MITEs of *Mutator* origin was roughly an order of magnitude lower in number and ranged from 592-3,626 elements. A fully autonomous *Mutator* element requires an intact transposase sequence. Among our *Mutator* TE set we identified very few intact transposase genes, from a low of 2 in *C. elegans* to a high or 18 in *C. briggsae*. Virtually all of the *Mutator* elements we annotated in *Caenorhabditis* genome sequences were partial fragments, non-autonomous and highly mutated (an example is shown for *C. elegans* in Fig. 8). Both *Mutator* elements and MITEs of *Mutator* origin had a high number of fragments 0-300bp in length. A *Mutator* sequence capable of transposition requires a pair of *∼*215bp terminal inverted repeats (TIRs) and a *∼*500bp transposase gene making a competent TE *>*1000bp in length. The abbreviated lengths and few intact transposase genes indicate the vast majority of *Mutator* elements are non-autonomous.

**Table 2.**
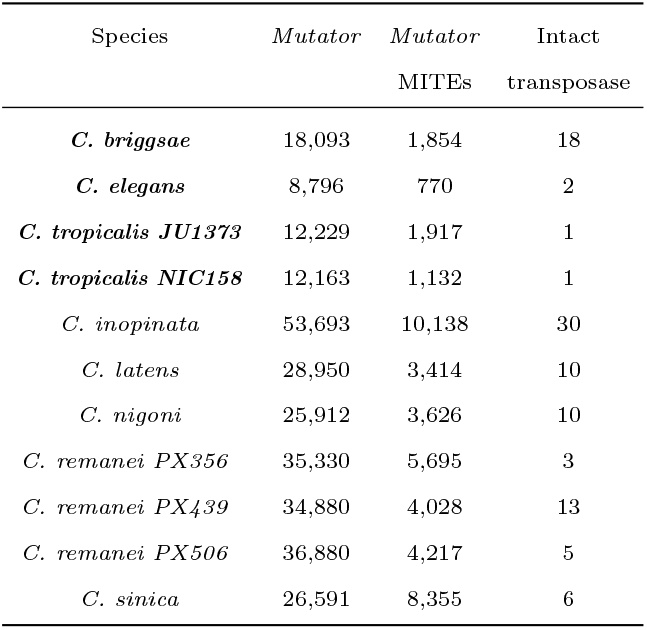
Numerical counts of *Mutator* -associated elements in *Caenorhabditis* genome sequences demonstrate uniformly high numbers of *Mutator* elements across the group and uniformly low numbers of intact transposase genes, indicating that virtually all of the annotated *Mutator* elements are either non-autonomous, partial fragments or both. The first column is *Caenorhabditis* species with strain name for species with multiple entries. The second column is total number of annotated *Mutator* elements, the third is MITEs that can be reliably associated to *Mutator* TEs through mutational patterns and the fourth is the number of intact transposase genes associated with the total set of *Mutator* elements. Self-fertile species are the first four rows in the table with species names in bold font.

**Fig. 8.**
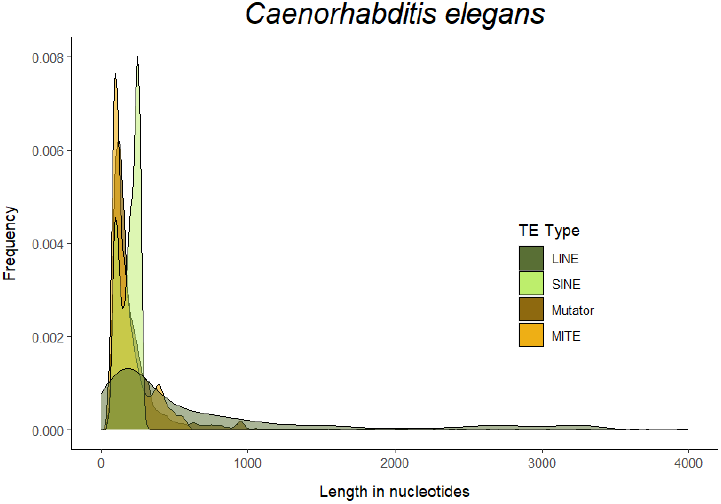
The majority of LINE, SINE, *Mutator* and *Mutator* -origin MITE sequences are truncated fragments of less than 500bp in length.

We further investigated autonomous and non-autonomous *Mutator* TEs by identifying those with competent TIRs (Table 3). Despite the high numbers of fragmented *Mutator* sequences in *Caenorhabditis* genomes (Table 2), very few have TIRs with sufficient sequence identity to competently transpose. *C. inopinata*, a close relative of the model *C. elegans* has an active tranposable element landscape potentially shaped by disruptive TE insertions into genes critical for the 26G small interfering (si)RNA gene silencing pathway (Kanzaki et al., 2018). The landscape of autonomous and non-autonomous *Mutator* TEs in *C. inopinata* shows that 95% of the non-autonomous *Mutator* TEs originate from just 5 of the 21 autonomous *Mutator* TEs. Phylogenetic reconstruction identified three groups of autonomous sequences (Supp. Fig. 6), potentially indicating three invasion events. One group contained a single autonomous sequence, one contained two sequences and the third group contained the remaining 18 sequences. Analysis of non-autonomous sequences with potentially competent TIRs showed that *>*98% of these sequences grouped with this large third group of autonomous sequences in phylogenetic reconstruction (Table 2; Supp. Fig. 7). Together, these patterns suggest most transposed *Mutator* sequences are rapidly mutated and quickly rendered incapable of transposition.

**Table 3.** Numerical counts of autonomous *Mutator* elements, non-autonomous *Mutator* elements with intact TIR sequences and non-autonomous Mutator elements with TIR sequences with 80% identity show that very few non-autonomous *Mutators* have the ability to transpose. The first column is *Caenorhabditis* species and strain name (self-fertile species are bolded), the second column is total number of intact autonomous *Mutator* elements, the third is non-autonomous *Mutators* with intact, competent TIR sequences and the fourth is non-autonomous *Mutator* elements with TIR sequences with 80% similarity to competent *Mutator* TIRs.

| Species | Autonomous<br>*Mutators* | Non-Aut.<br>Exact TIRs | Non-Aut.<br>80% TIRs |
| --- | --- | --- | --- |
| ***C. briggsae*** | 8 | 0 | 19 |
| ***C. elegans*** | 2 | 0 | 0 |
| ***C. tropicalis JU1373*** | 1 | 13 | 13 |
| ***C. tropicalis NIC158*** | 1 | 0 | 0 |
| *C. inopinata* | 21 | 49 | 556 |
| *C. latens* | 7 | 0 | 4 |
| *C. nigoni* | 6 | 0 | 6 |
| *C. remanei PX356* | 3 | 0 | 0 |
| *C. remanei PX439* | 6 | 0 | 28 |
| *C. remanei PX506* | 5 | 3 | 50 |
| *C. sinica* | 3 | 0 | 2 |

**Table 4.**
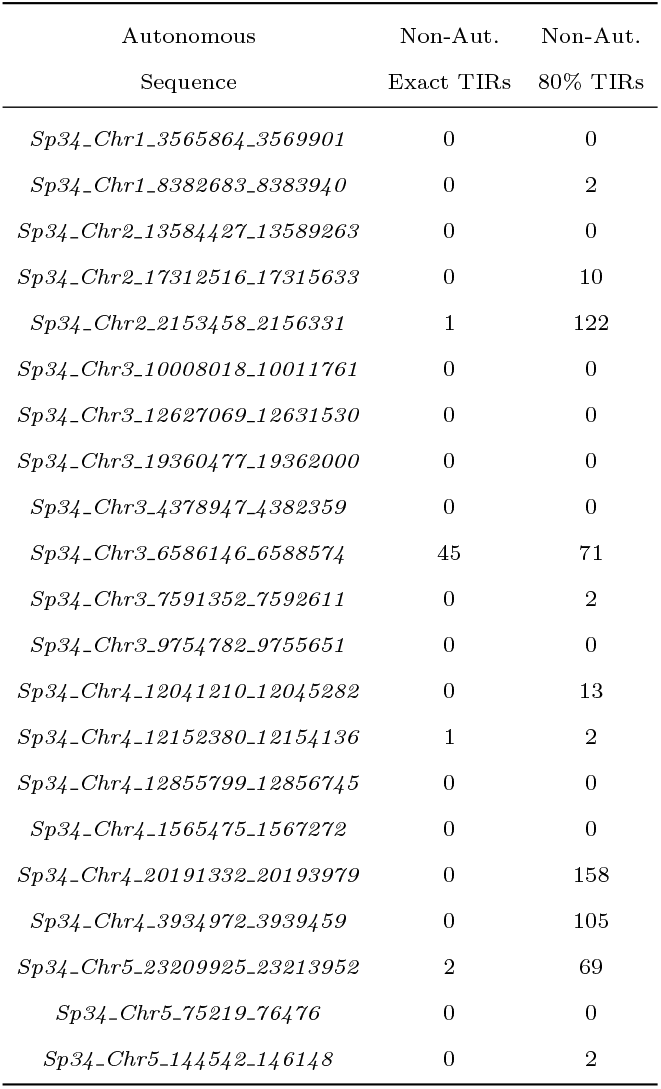
Numerical counts of autonomous *Mutator* elements, non-autonomous *Mutator* elements with intact TIR sequences and non-autonomous Mutator elements with TIR sequences with 80% identity in *C. inopinata* show that few autonomous *Mutators* produced 94.4% of the non-autonomous elements with recognizable TIRs. The first column is *C. inopinata* autonomous *Mutator sequence*, the second column is non-autonomous *Mutators* with intact, competent TIR sequences and the fourth is non-autonomous *Mutator* elements with TIR sequences with 80% similarity to competent *Mutator* TIRs.

## Discussion

The question of how and under what conditions transposable elements might self-regulate has a developed history in evolutionary biology (Charlesworth and Charlesworth, 1983; Charlesworth and Langley, 1986; Langley et al., 1983). Previous theoretical studies have shown stable persistence of transposable elements under narrow sets of conditions and created an apparent paradox with the high transposon content found in most eukaryotic genomes (Kelleher et al., 2020). Here, we have presented deterministic and stochastic models that demonstrate regulatory logic can result in long-term stable persistence of transposable elements. We used our model results to structure investigations of autonomous and non-autonomous elements in *Caenorhabditis* genome sequences and found support for both competitive and parasitic regulatory interactions. Together, our theoretical and comparative genomic results suggest broad variation in regulatory logic across TE families.

### Theoretical studies of intrinsic transposable element regulation

Early theoretical studies of transposable elements were fascinated with ideas of self-attenuation and self-regulation (Charlesworth and Charlesworth, 1983; Charlesworth and Langley, 1986; Langley et al., 1983). Without some mechanism of containment, transposable elements could conceivably proliferate until they took over entire genomes or decreased fitness to the point of population extinction. Mapping out the evolutionary processes that could support the broad diversity of TEs found in empirical populations was an important theoretical problem (Kelleher et al., 2020).

Despite extensive theoretical study, the conditions under which transposable elements were predicted to stably persist in populations were fairly narrow. Charlesworth and Charlesworth (1983) found that TEs could be stably maintained with either synergistic interactions for fitness, where the decrease in fitness accumulates faster than linearly with increasing numbers of transposable elements, self-regulation of transposition or both. Self-regulation of transposition was only achieved with unrealistic modeling assumptions including the total absence of selection (Charlesworth and Charlesworth, 1983), free recombination and no linkage (Langley et al., 1983), or highly restrictive rates of recombination (Charlesworth and Langley, 1986). Similarly, empirical studies have found that synergistic epistasis occurs (Lee, 2022) but is not ubiquitous (Maisnier-Patin et al., 2005; de Visser et al., 2011).

More recent theoretical treatments increased the space under which transposable elements may self-regulate by modeling autonomous and non-autonomous elements (Le Rouzic and Capy, 2006; Omole and Czuppon, 2024), similar to the models we have developed here. However, these studies still predicted short-term persistence of TEs in the population or required evolution in the absence of selection and fitness effects.

Despite broad empirical patterns of transposable element stability, the predicted space for self-attenuating transposition has remained quite small over 40 years of study. This has led some authors to propose the ‘case closed’ on the potential for transposable elements to engage in any meaningful self-regulation (Kelleher et al., 2020).

A major contribution of our results is to demonstrate the theoretical potential for long-term, stable maintenance of transposable elements. Our simulated populations evolving with parasitic regulatory interactions between autonomous and non-autonomous transposable elements showed stable persistence of TEs across a broad range of population genetic factors including varying recombination, mutation and outcrossing rates, and population sizes. Our results suggest that regulatory logic may play an important, determining role in the evolution and stability of transposable elements in eukaryotic genomes.

### The maintenance of transposable elements in selfing and asexual populations

Theoretical studies have uniformly predicted little space for transposable element maintenance in asexual or selfing populations (Hickey, 1982; Charlesworth and Charlesworth, 1983; Boutin et al., 2012). Our study largely agrees with these findings and in our models transposable elements were only maintained in partially selfing populations where at least 50% of the individuals reproduced through outcrossing. However, these theoretical results are at odds with empirical studies documenting abundant transposable elements in the genomes of selfing and asexual organisms (Nowell et al., 2021; Fierst et al., 2015; Adams et al., 2023; Bonchev and Willi, 2018; Mattila et al., 2020). How can these two phenomenon-be reconciled?

Importantly, theoretical studies and population models predicting an erosion of transposable elements in asexual and selfing organisms still envision the loss of transposable elements as a process occurring over time. Empirical studies have found mixed evidence for this in experimental evolution studies (Bast et al., 2019; Chen and Zhang, 2021) but found the process dependent on the dynamics of the population, elements and genomic structure (Henault et al., 2024). At the same time, organisms are constantly subject to TE invasions (Panaud, 2016; Peccoud et al., 2017; Guan et al., 2022; Paulat et al., 2023). Transposable element families in asexual and selfing populations may have fundamentally different dynamics from those in sexual populations, with rapid rates of turnover resulting from continual cycles of invasion and loss. An interesting avenue of future research would be phylogenetic comparative analysis of TEs across related sexual and asexual or selfing species, searching for ephemeral dynamics or differing rates across evolutionary time.

### The evolution of extrinsic transposable element regulation

We focused on the potential for internal or intrinsic regulatory interactions between and within transposable elements to stabilize TEs in populations. In recent years external or extrinsic TE regulation has been an exciting field of study (Brennecke et al., 2007). Small RNAs, including microRNAs (Oliver et al., 2022), small interfering (si-)RNAs (Wei et al., 2014) and PIWI-interacting or piRNAs (Ozata et al., 2018), target TE sequences for degradation and are an important mechanism of extrinsic regulation. In our study populations could go to extinction due to TE proliferation. This is likely not a reasonable scenario and in natural populations continually heavy TE burdens may facilitate the evolution of genomic resistance mechanisms, including extrinsic regulation capable of slowing proliferation.

Small RNA regulation systems play vital roles in many organisms but are not a universal mechanism of TE containment and regulation. The systems are absent in some basal eukaryotes (Ryan et al., 2013; Maxwell et al., 2012) and evolutionary reconstruction suggests parallel loss in others (Mondal et al., 2018; Fontenla et al., 2021; Sarkies, 2024; Sarkies et al., 2015). An interesting question for future study is if these organisms have fundamentally different patterns of genomic TEs from those with developed small RNA systems targeting and regulating TEs. Developing genomic data could facilitate comparative studies investigating how the number, diversity and location of transposable element sequences and families differs between organisms with and without small RNA systems of regulation.

### Empirical studies of autonomous and non-autonomous transposable elements

Empirical studies of autonomous and non-autonomous TEs have reported interesting dynamics. For example, Robillard et al. (2016) tracked the novel introduction of autonomous and non-autonomous *Mariner* transposons into *Drosophila melanogaster* populations. They found that rapid proliferation of the non-autonomous element led to extinction of both types of elements within 100 generations. In contrast to this rapid extinction, autonomous -non-autonomous partners also appear capable of coexisting and coevolving for long periods (Tanskanen et al., 2007; Wicker et al., 2022). Together, these studies suggest that autonomous -non-autonomous regulatory logic can determine the stability or instability of transposable elements in genomes.

Hatanaka et al. (2024) studied transposons from the *hAT* superfamily that have inserted into the genome of *C. inopinata*, a sister species of *C. elegans*. The *C. elegans* genome lacks *hAT* elements (Woodruff and Teterina, 2020), suggesting a relatively recent insertion event. Active transposable elements in *C. inopinata* have inserted into essential genes in the 26G si-RNA pathway (Kanzaki et al., 2018) and *C. inopinata* and its sister species *C. elegans* appear to have diverged in genomic TE landscape (Woodruff and Teterina, 2020) and TE-associated gene expression (Kawahara et al., 2023). Similar to our comparative genomic results Hatanaka et al. (2024) identified a biased pattern of autonomous -non-autonomous elements (Table 2) with just 4 autonomous *hAT* elements and over 1,000 deletion-related non-autonomous miniature inverted-repeat transposable elements (MITEs). The genomic location of TEs can play important roles in recombination and mutation with differing dynamics and gene density in chromosome arms and centers in *Caenorhabditis* (Rockman and Kruglyak, 2009). The genomic distribution of *hAT* -associated MITEs and other highly abundant non-autonomous elements can influence evolution of the genome and have consequences for multiple dimensions of phenotypic evolution.

### The dynamics of transposable element invasion

Our empirical investigation of autonomous and non-autonomous *Mutator* elements in *Caenorhabditis* discovered a ubiquitous pattern of fragmented sequences and mutated TIRs rendering non-autonomous elements incapable of transposition and essentially ‘silencing’ them in the genome. The course of a transposable element invasion into a genome can be likened to organismal infection with a novel pathogen. The early, initial stages of infection are characterized by dynamic proliferation, here tranposition within the genome (Robillard et al., 2016). Within a period of time the infection or invasion is suppressed, here by mutations rendering the sequences incompetent for transposition. After the initial period of dynamic proliferation, TE-associated sequences remain in the genome although inactive and not participating in, or governed by, regulatory logic such as Competitive or Parasitic dynamics. Importantly, these silenced sequences can now create pools of novel variation for regulation (Du et al., 2024) and *de novo* gene creation (Peng and Zhao, 2024). Regulatory logic may play a critical role in the acute stages of transposable element invasion into a genome and lose importance as the invasion shifts into a chronic, silenced state.

## Conclusion

Transposable elements constitute large portions of eukaryotic genomes with diverse numbers and types of elements. Despite this, there is little understanding of the evolutionary processes and population genetic factors that produce patterns of TE diversity. We have presented deterministic and stochastic theory studying the role that regulatory logic can play in determining the stability or instability of systems of autonomous and non-autonomous transposons. Our results show that regulatory interactions can interact with population genetic factors to result in highly stable systems of transposable elements. We investigated our predictions in *Caenorhabditis* genome sequences and found that empirical patterns of autonomous -non-autonomous TEs vary broadly. Rapid mutational decay likely renders non-autonomous TEs incapable of transposition and suggests that TE regulatory logic is relevant for early, acute stages of TE invasion. Together, our results demonstrate that regulatory logic can stabilize TEs and that empirical TEs employ diverse regulatory strategies.

## Author contributions statement

Must include all authors, identified by initials, for example: J.L.F. conceived the research(s), V.K.E. and J.L.F. conducted the analyses(s), wrote and reviewed the manuscript.

## Acknowledgments

The authors thank the members of the Fierst lab for helpful feedback and critical discussion of the research and manuscript.

## Data availability

Code and output associated with the deterministic and stochastic models are available at https://github.com/jannafierst/TEParasites. Bioinformatic scripts and workflows are available at https://github.com/vkeggers/Caenorhabditis_TE_autonomousANDnonautonomous.

*Caenorhabditis* genome sequences are available at NCBI under the accessions:

*C. remanei* PX356 Bioproject PRJNA248909

*C. remanei* PX439 Bioproject PRJNA248911

*C. remanei* PX506 Bioproject PRJNA577507

*C. latens* PX534 Bioproject PRJNA248912

*C. briggsae* AF16 Bioproject PRJNA20855

*C. nigoni* JU1422 Bioproject PRJNA384657

*C. elegans* N2 Bioproject PRJNA158, PRJNA13758

*C. tropicalis* NIC58, JU1373 Bioproject PRJNA662844

*C. inopinata* NKZ35 Bioproject PRJDB5687

*C. sinica* ZZY0401 Bioproject PRJNA194557

## Supplementary Figures

**Supplementary Figure 1.**
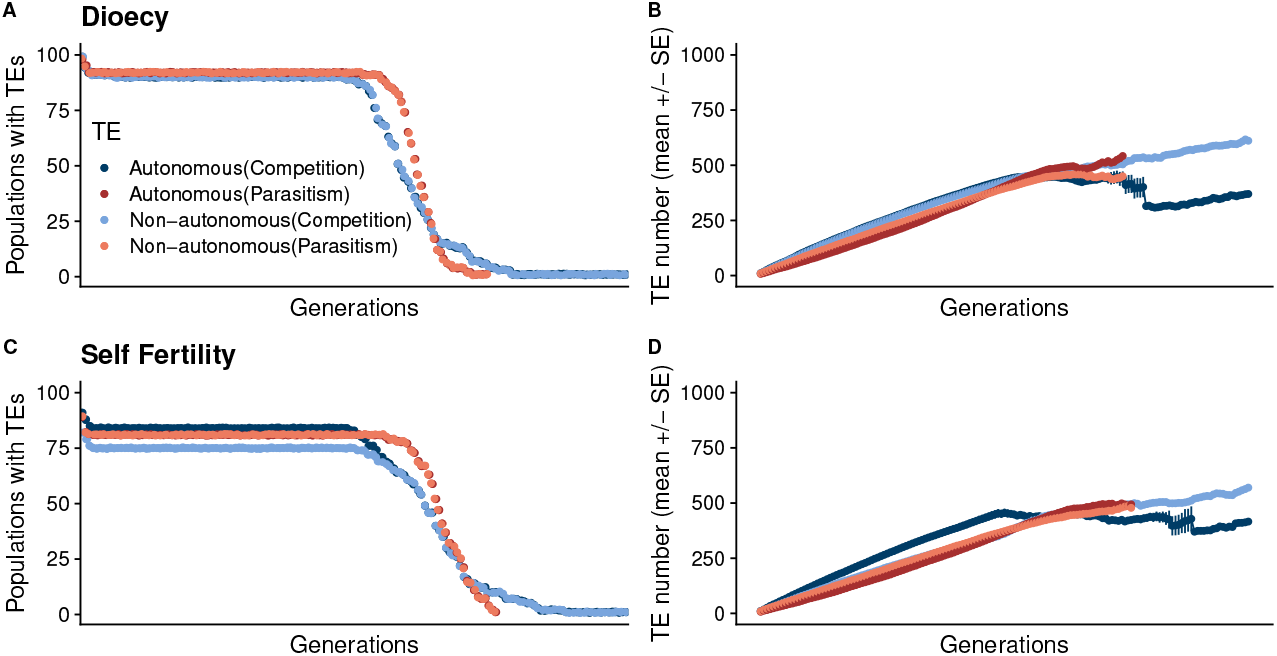
In small (here, N=100 individuals) dioecious populations both (A) Competitive and Parasitic TE regulation resulted in one of two trajectories, either early TE loss or TE proliferation to the point of extreme fitness loss and population crash. The populations were evolved for 500,000 generations and all simulations started with 100 populations. (B) The mean number of TEs (+/-SE) were calculated for populations with TEs present. C) In small self-fertile populations (here, N=100 individuals) both Competitive and Parasitic TE regulation resulted in similar trajectories of early TE loss or TE proliferation, fitness loss and population crash. D) TEs could proliferate until the population crashed

**Supplementary Figure 2.**
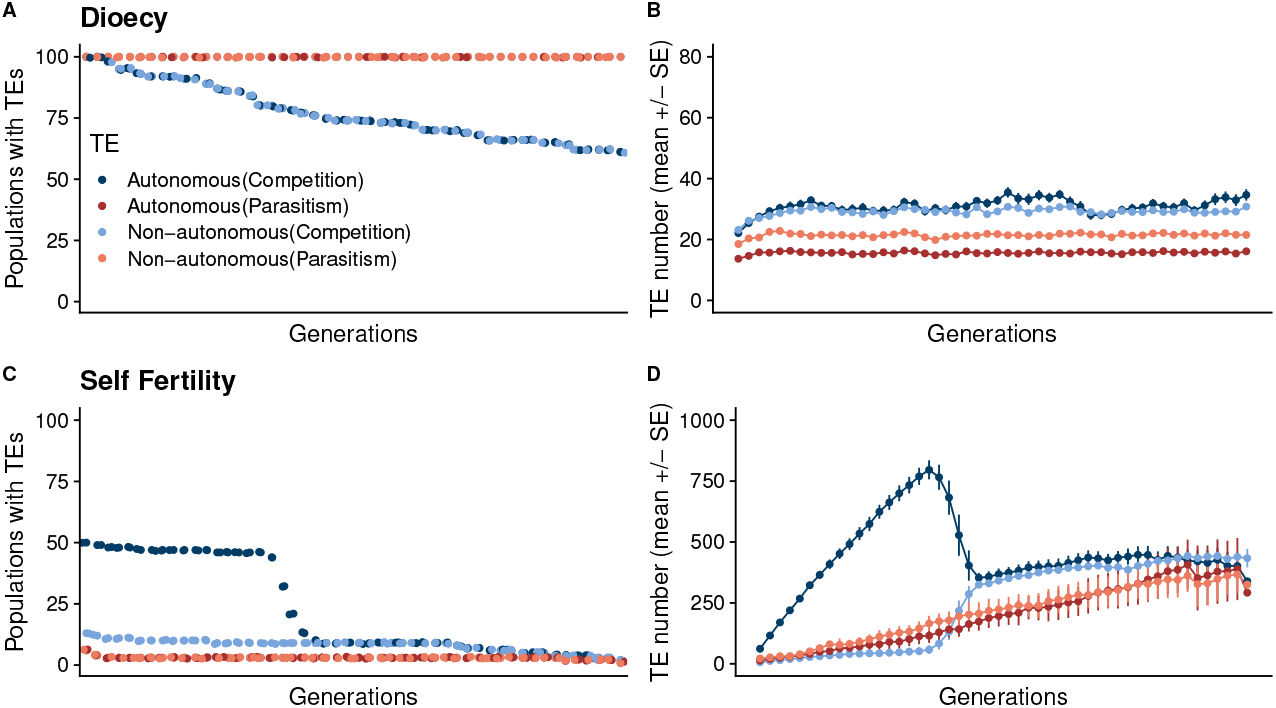
Increasing the recombination rate to 1e-2 did not alter the qualitative trajectory of TEs in the simulated populations. At large size (here, N=1000) dioecious populations A) evolving with Parasitic TE regulation achieved Stable Maintenance while populations evolving with Competitive TE regulation did not. B) For those populations that did retain TEs, the mean abundance was lower when compared with populations evolving with lower recombination rates. C) Self-fertile populations showed the same trajectories of TE proliferation and loss when evolving with high recombination rates including D) decoupling of the autonomous and non-autonomous TEs.

**Supplementary Figure 3.**
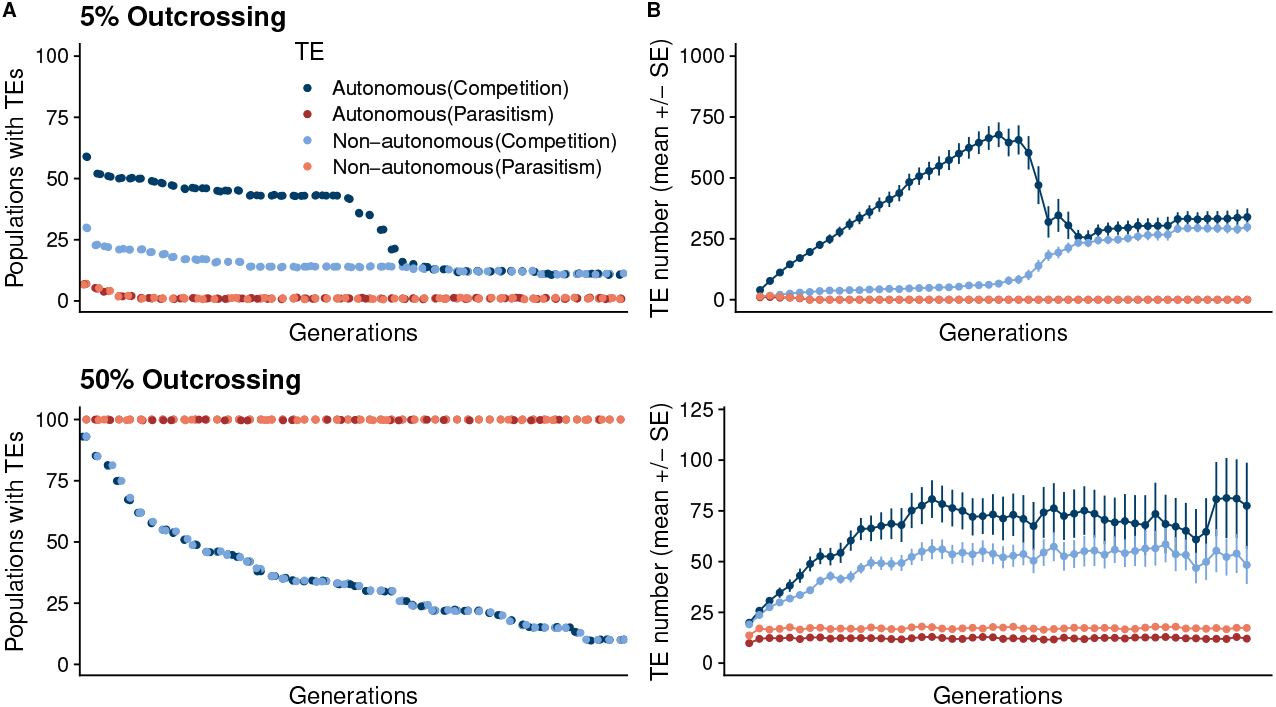
Populations evolving with both Competitive and Parasitic TE regulation follow the trajectories of fully self-fertile populations at A,B) 5% outcrossing and fully outcrossing populations at C,D) 50% outcrossing.

**Supplementary Figure 4.**
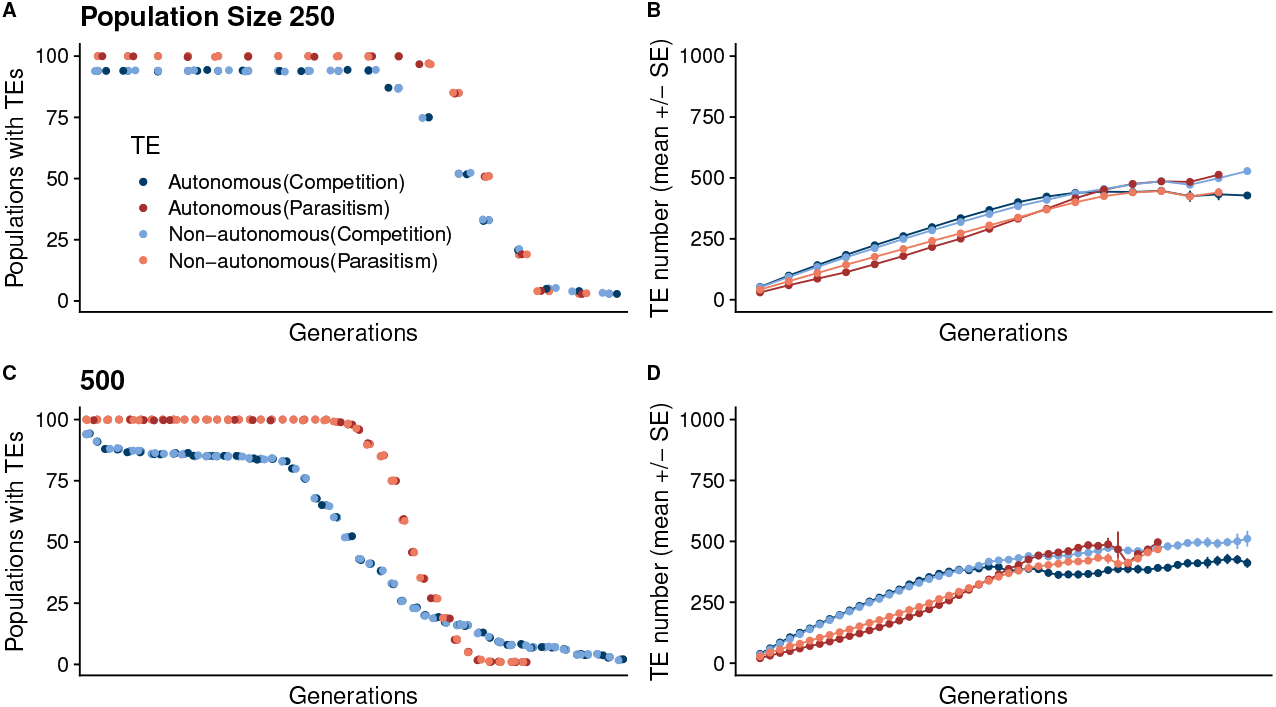
Small populations of A,B) 250 and C,D) 500 individuals show similar patterns to populations evolving with 100 individuals (SFigure 1). TEs follow either the ‘Early TE Loss’ or ‘TE Proliferation & Population Crash’ trajectories.

**Supplementary Figure 5.**
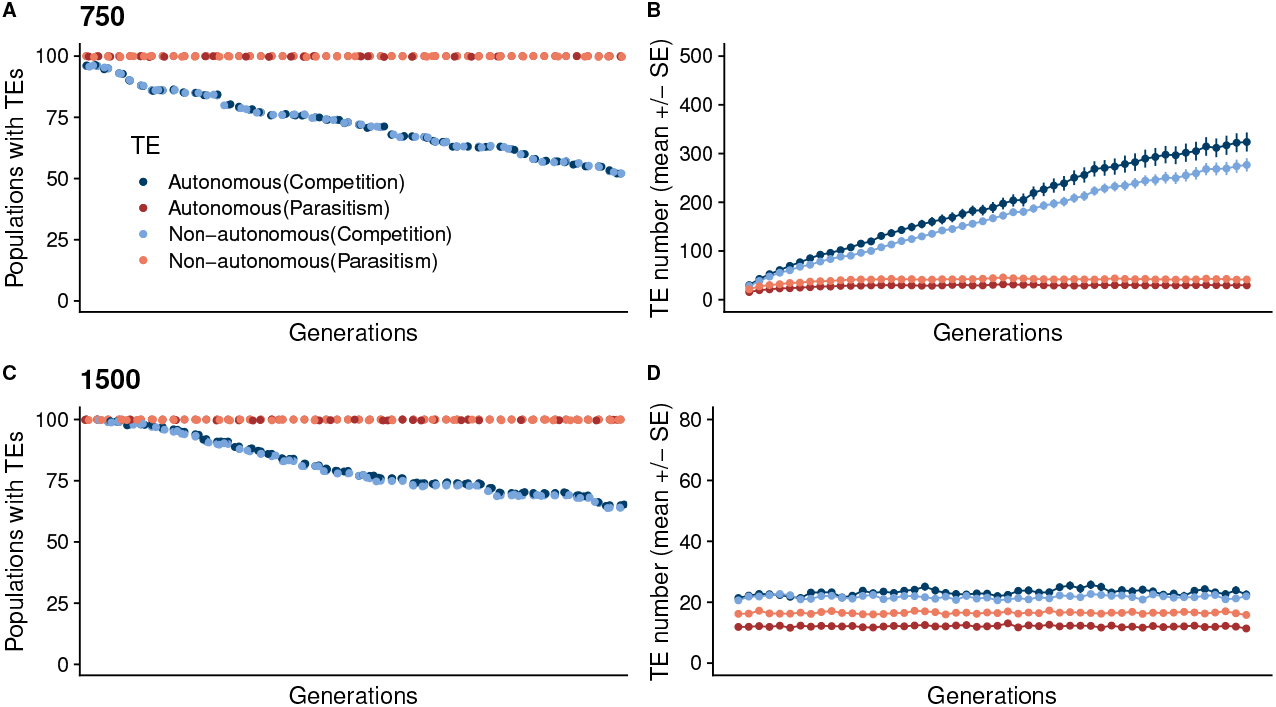
Populations evolving with larger population sizes show similar TE patterns to populations evolving with 1000 individuals (Fig. 5) Parasitic TE regulation achieve the stable maintenance trajectory at population sizes of A-B) 750 and C-D) 1500 individuals. With Competitive regulation TEs are maintained for longer periods but lost according to the ‘Eventual TE Loss’ trajectory. In populations of D) 1500 individuals TEs are maintained at lower mean levels in populations evolving with 750 individuals.

**Supplementary Figure 6.**
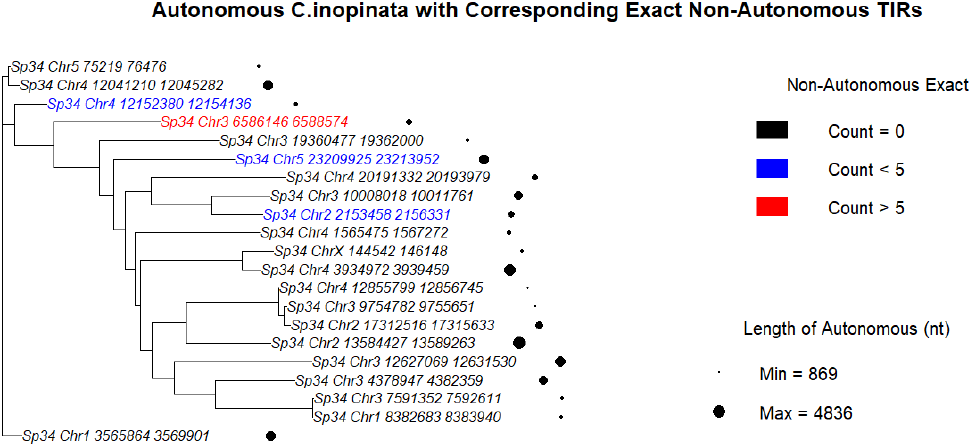
Phylogenetic reconstruction of autonomous *Mutator* sequences in *C. inopinata* (also known as *Caenorhabditis* Species 34) shows the transposons form 3 clades, potentially reflecting 3 separate *Mutator* invasions. One clade contains a single sequence, one contains two sequences and the third clade contains the remaining 18 sequences. Of these, 17 have no corresponding non-autonomous elements with intact TIRs, three have *<*5 corresponding non-autonomous elements with intact TIRs and the remaining one autonomous element has *>*5 corresponding non-autonomous elements.

**Supplementary Figure 7.**
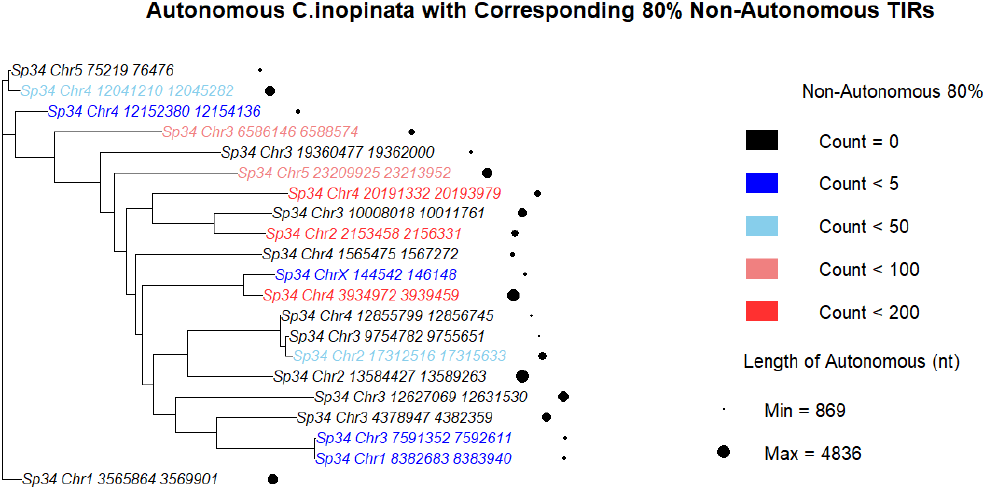
Phylogenetic reconstruction of autonomous *Mutator* sequences in *C. inopinata* shows that 7 autonomous sequences each retain *>*80% TIR identity with *>*50 non-autonomous sequences. The remaining 14 autonomous sequences retain *>*80% identity with 5 or fewer non-autonomous *Mutator* elements.

